# HOXDeRNA activates a cancerous transcription program and super-enhancers genome-wide

**DOI:** 10.1101/2023.06.30.547275

**Authors:** Evgeny Deforzh, Prakash Kharel, Anton Karelin, Pavel Ivanov, Anna M. Krichevsky

## Abstract

**Background:** The origin and genesis of highly malignant and heterogenous glioblastoma brain tumors remain unknown. We previously identified an enhancer-associated long non-coding RNA, LINC01116 (named HOXDeRNA here), that is absent in the normal brain but is commonly expressed in malignant glioma. HOXDeRNA has a unique capacity to transform human astrocytes into glioma-like cells. This work aimed to investigate molecular events underlying the genome-wide function of this lncRNA in glial cell fate and transformation.

**Results:** Using a combination of RNA-Seq, ChIRP-Seq, and ChIP-Seq, we now demonstrate that HOXDeRNA binds *in trans* to the promoters of genes encoding 44 glioma-specific transcription factors distributed throughout the genome and derepresses them by removing the Polycomb repressive complex 2 (PRC2). Among the activated transcription factors are the core neurodevelopmental regulators SOX2, OLIG2, POU3F2, and SALL2. This process requires an RNA quadruplex structure of HOXDeRNA that interacts with EZH2. Moreover, HOXDeRNA-induced astrocyte transformation is accompanied by the activation of multiple oncogenes such as EGFR, PDGFR, BRAF, and miR-21, and glioma-specific super-enhancers enriched for binding sites of glioma master transcription factors SOX2 and OLIG2.

**Conclusions:** Our results demonstrate that HOXDeRNA overrides PRC2 repression of glioma core regulatory circuitry with RNA quadruplex structure. These findings help reconstruct the sequence of events underlying the process of astrocyte transformation and suggest a driving role for HOXDeRNA and a unifying RNA-dependent mechanism of gliomagenesis.

## Background

Glioblastoma (GBM, or grade IV astrocytoma) is the most prevalent malignant primary tumor of the central nervous system in adults, with a median survival of 15 months. Despite substantial research, molecular events underlying the transformation of normal cells to glioma-initiating cells and the development of this highly heterogeneous disease are poorly understood. Glioma master transcription factors (TFs) and other genes responsible for glioma cell identity are predominantly silenced in brain astrocytes, the cells of origin of glioma [1]. The silent status of these genes depends on the Polycomb repressive complex 2 (PRC2) component EZH2, which catalyzes the tri-methylation of lysine 27 on histone H3 (H3K27Me3) [2]. EZH2 also binds multiple RNA species *in vitro* and *in vivo* [3–6]. Although the selectivity of EZH2-RNA binding is still debatable, essential regulatory functions of several lncRNAs, such as Xist and HOTAIR [7, 8], were linked to their PRC2-binding capacity. In line with the promiscuous RNA binding of PRC2, recent reports also suggested EZH2’s high affinity and specificity for G-rich sequences and RNA quadruplex structures (rG4) [9, 10]. However, the biological significance of lncRNA-EZH2 interactions, and particularly of rG4-EZH2 binding, remains to be elucidated.

We have previously shown that the glioma-specific enhancer RNA (eRNA) LINC01116 (named HOXDeRNA here), not expressed in the normal brain but commonly induced in glioma, activates the transcription of all HOXD genes in GBM [11]. The underlying mechanism involves CTCF/cohesin-dependent looping between the HOXD locus and the HOXDeRNA-encoding enhancer located 500 kb downstream. In this report, we show that HOXDeRNA derepresses key glioma genes not only *in cis* but also *in trans* in a genome-wide manner. We describe a decoy mechanism, which involves rG4-dependent recruitment of HOXDeRNA to PRC2-covered transcription start sites (TSS) of glioma driver genes, including master TFs, followed by the removal of PRC2 from the gene bodies. Targeted base editing of a specific rG4 in HOXDeRNA abrogates HOXDeRNA recruitment to chromatin and the removal of PRC2, repressing the glioma signature genes and transformation.

## Results

### Activation of HOXDeRNA in astrocytes induces glioma transcriptional programs

We confirmed and expanded prior findings [11] that HOXDeRNA was not expressed in normal brain tissues, astrocytes, oligodendrocytes, and neural stem cells but was actively transcribed in glioma and GBM tumor tissues, cells, and glioma stem cells (GSCs) using multiple datasets (Supplemental Figure 1A and B). To investigate HOXDeRNA role in glioma biology, we employed human immortalized astrocytes [12–14] which transcriptome mirrors that of primary cortical astrocytes (Supplemental Figure 1C). We showed previously that activation of HOXDeRNA with CRISPR activation system promoted the transformation of astrocytes into glioma-like spheroids [11]. Differential gene expression analysis of control astrocytes and the astrocytes transformed by overexpressing HOXDeRNA revealed 2698 activated and 3456 repressed genes in the transformed spheroids (FC> 2, p-val <0.01) (Figure 1A, B). Gene ontology (GO) analysis demonstrated that expressions of cell-cell junction, focal and cell adhesion genes, including cadherins, claudins and integrins, decreased after the astrocyte transformation, suggesting a molecular basis for the observed phenotypic transition from adherent cells to spheroids (Figure 1C). Notably, the genes upregulated in transformed cells were related to transcriptional regulation, growth factors, and tyrosine kinases, which were known to be associated with glioma biology. Among the genes upregulated by HOXDeRNA activation were those encoding 44 glioma-specific master TFs, including SOX2, OLIG1, OLIG2, HEY2 [1, 15]. In addition, expressions of multiple key factors frequently mutated and/or activated in gliomas, including the established therapeutic targets such as EGFR, PDGFRA, BRAF, and TERT, were upregulated by HOXDeRNA (Figure 1B). Both groups of genes define transcriptional programs and glioma-like identity of the transformed astrocytes. The machine learning model trained on 30 differentially expressed genes demonstrated the respective association of control and transformed astrocytes with the normal brain and glioma, based on their transcriptomics (Figure 1D).

**Figure 1.**
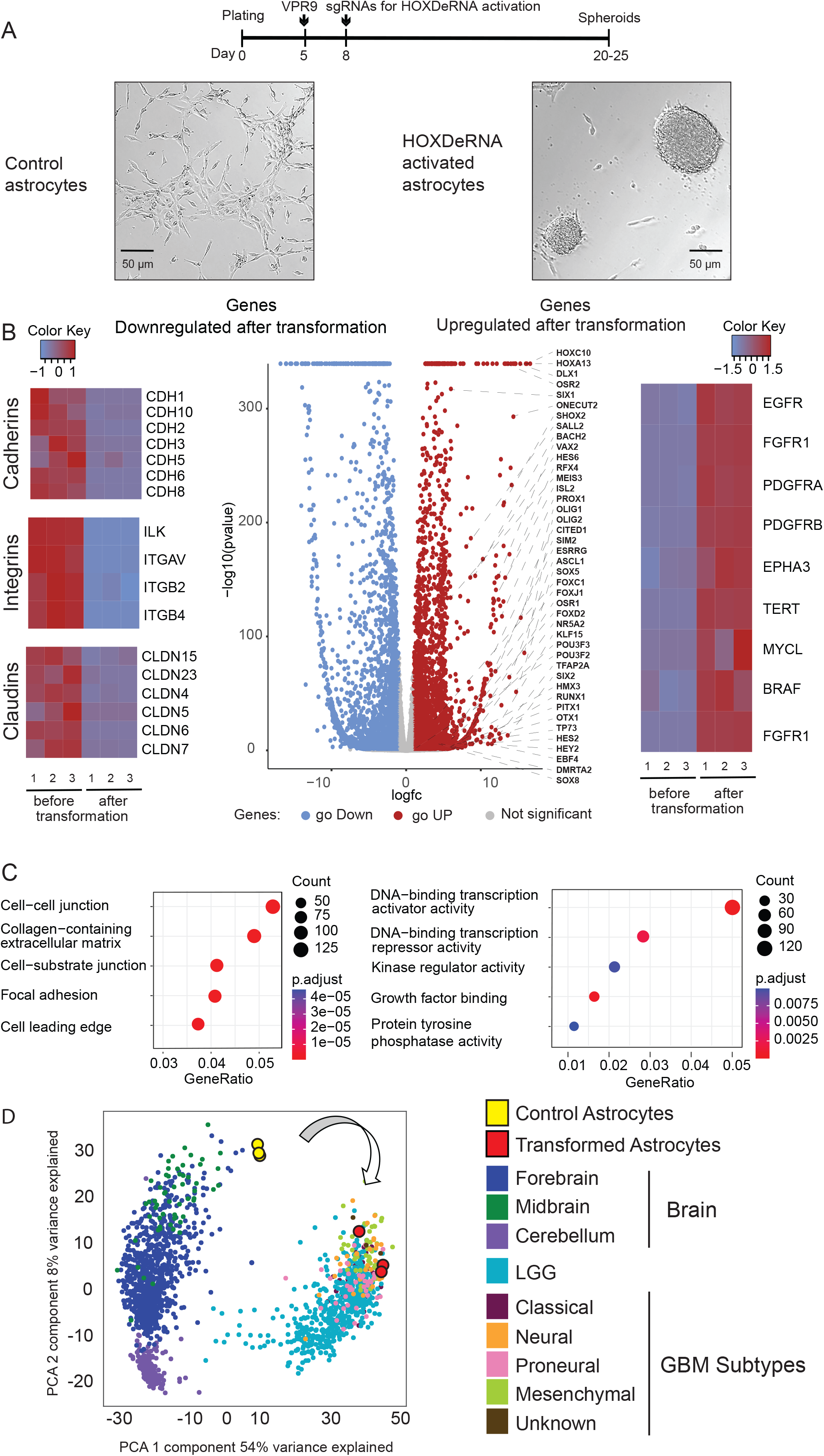
Targeted activation of the HOXDeRNA leads to astrocyte transformation with phenotypic and transcriptomic switch to glioma. A. Timeline for transduction of astrocytes with the CRISPR activation system leading to transformation (top). Representative images of astrocytes transduced with non-targeting sgRNA (control astrocytes) and HOXDeRNA-activating sgRNA (HOXDeRNA activated astrocytes) are shown. B. Volcano plot (middle) showing differentially expressed genes (DEG) between the non-transformed and transformed astrocytes. The genes upregulated in transformed astrocytes (red), downregulated in transformed astrocytes (blue) (fold change > 2 and adjusted p-value 0.01), and 10% of non-significantly changed genes (grey) are shown. The 44 glioma specific TFs that are upregulated after astrocyte transformation are indicated. The heatmaps exhibit z-scores for cell junction and cell adhesion genes downregulated in transformed astrocytes (left, n=3) and major glioma-associated genes upregulated in transformed astrocytes (right, n=3). C. The top 5 categories of GO gene sets downregulated (left) or upregulated (right) in transformed astrocytes shown for DEG (FC>2, p<0.01). D. Сontrol and transformed astrocytes are associated with “normal brain” and “glioma” expression signatures, respectively. Machine learning model trained on forebrain (n=857 samples), cerebellum (n=214), midbrain (n=57), low grade glioma (LGG, n=522), as well as mesenchymal (n=54), classical (n=41), proneural (n=39), neural (n=28), and unknown subtype (n=4) GBM samples from TCGA and GTEX datasets classifies control and transformed astrocytes as “normal brain” and “glioma”, correspondingly, based on the RNA-Seq data. PCA visualization is shown, and every dot represents an individual sample (see Methods for details).

### HOXDeRNA binds and removes PRC2 from genes encoding glioma-specific master transcription factors in transformed astrocytes

The RNA-Seq data showing that HOXDeRNA activation changed the expression of thousands of genes suggested a genome-wide regulatory activity of HOXDeRNA. To investigate this activity, we employed the Chromatin Isolation by RNA Purification coupled with sequencing (ChIRP-seq) technique, which captures direct interactions between the RNA of interest and chromatin. We detected 1085 common chromatin binding sites for HOXDeRNA in transformed astrocytes and three GSC lines (Figure 2A, B). Genomic distributions of HOXDeRNA binding sites in transformed astrocytes and GSCs were similar, with 61-65% binding events observed in gene promoters and 25-28% in the intergenic regions, and were mapped to all chromosomes (Supplemental Figure 2A), suggesting a global role for HOXDeRNA in gene transcription. Notably, HOXDeRNA bound almost exclusively to the promoters of genes that were upregulated in transformed astrocytes, but not the genes whose expression was repressed (Figure 2C, D; Supplemental Figure 2B, C). The corresponding gene lists (Supplemental Table 1) were obtained from the differential gene expression analysis shown in Figure 1B (volcano plot).

**Figure 2.**
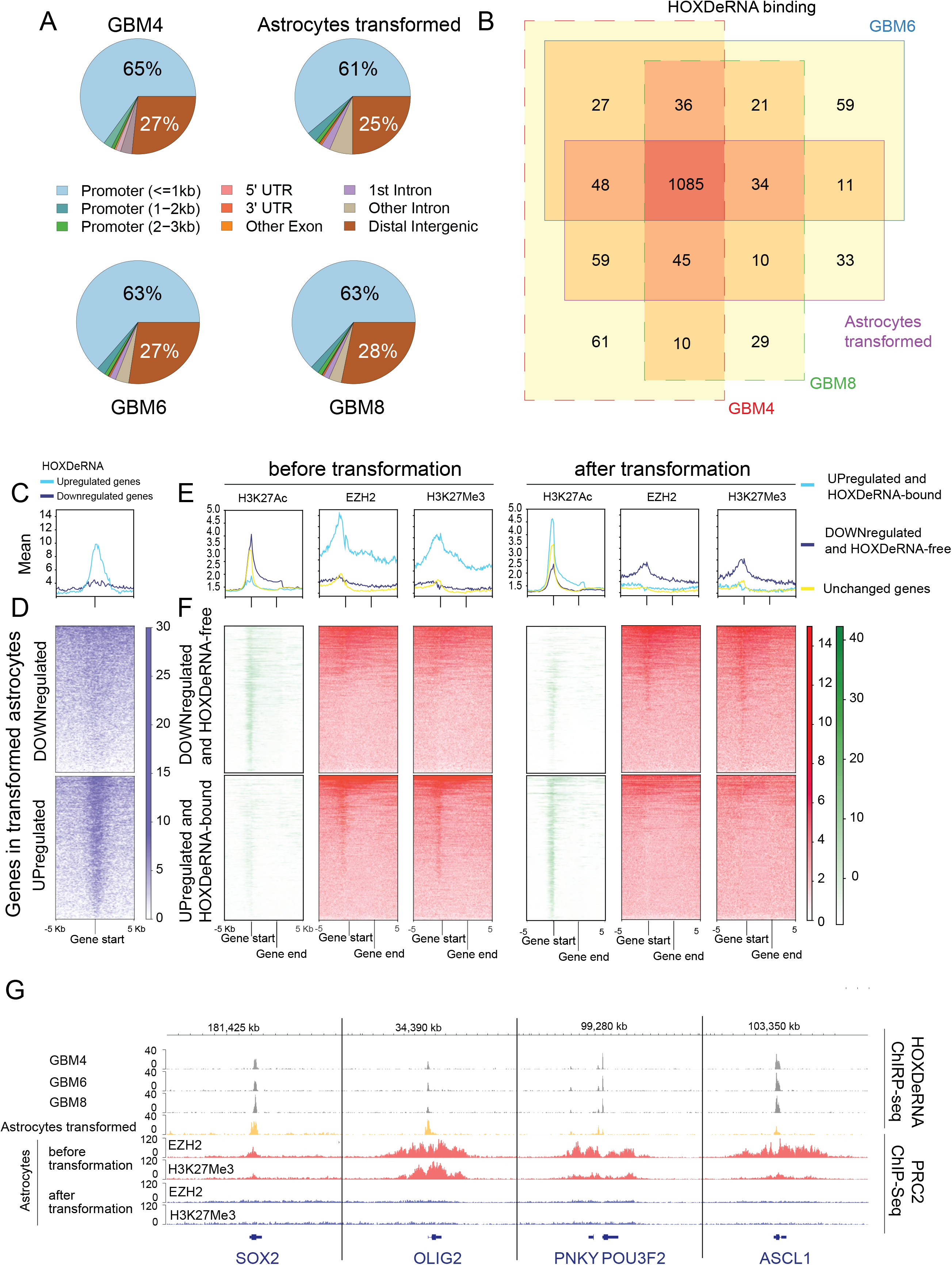
Genome-wide binding of HOXDeRNA is associated with exclusive PRC2 removal from transformation-induced genes. A. ChIRP-seq analysis demonstrates that transformed astrocytes and three GSC lines (GBM4, GBM6 and GBM8) exhibit similar distribution of HOXDeRNA peaks across the genome, with most peaks mapped to gene promoters. B. HOXDeRNA binds to the same gene promoters in transformed astrocytes and GSCs. HOXDeRNA ChIRP peaks were annotated to the nearest genes, and the gene lists produced for the four cell types were intersected and visualized as a Venn diagram (see Methods for details). C, D. ChIRP-seq raw read coverage signal, representing the HOXDeRNA binding at the TSS (+/- 5Kb) of the forward strand of genes downregulated or upregulated after astrocyte transformation, is visualized as an average value for each group (C) or for individual genes (D). The heatmap is accompanied by a colour scheme representing the value of the raw read counts. A similar coverage for reverse strand genes is shown in Supplemental Figure 2B, C. E, F. Epigenetic status of genes upregulated or downregulated after astrocyte transformation. H3K27Ac, H3K27Me3, and EZH2 ChIP-seq raw signals covering gene bodies of the positive strand (+/- 5Kb) were normalized to gene length and counted, followed by visualization of the average signal for 3 groups: genes upregulated after astrocytes transformation and bound by HOXDeRNA, downregulated after astrocyte transformation and HOXDeRNA-free, and unchanged genes (E). Individual gene body coverage values were visualized as heatmaps (F) (see reverse strand gene coverage in Supplemental Figure 2D, E). G. HOXDeRNA ChIRP-seq tracks in GSCs and transformed astrocytes, aligned with PRC2 (H3K27Me3, EZH2) ChIP-seq coverage, before and after astrocyte transformation, visualized for selected glioma master TF genes.

We next investigated the epigenetic status of HOXDeRNA-occupied genes in astrocytes, using H3K27Ac as the mark of active chromatin state and H3K27Me3 and EZH2 (PRC2) as the marks associated with repressed chromatin. Notably, genes bound by HOXDeRNA and upregulated after astrocyte transformation were occupied by PRC2 and depleted of the H3K27Ac mark before transformation and significantly decreased their PRC2 coverage after transformation (Figure 2E, F, Supplemental Figure 2D, E). This contrasted with the unchanged and downregulated genes not bound by HOXDeRNA (Figure 2E, F, Supplemental Figure 2D, E). Importantly, all genes encoding the 44 glioma-specific master TFs were bound by HOXDeRNA and lost their PRC2 marks after cell transformation (Figure 2G, Supplemental Figures 2F, G, H, I and Supplemental Figure 3). We highlight this remarkable effect using representative master TFs SOX2, OLIG2, POU3F2, and ASCL1, their genes were bound by HOXDeRNA in three GSCs and transformed astrocytes (Figure 2G, grey and yellow tracks) and HOXDeRNA binding in transformed astrocytes was associated with the reduced PRC2 repression (Figure 2G, blue and red tracks, respectively; Supplemental Figure 2J). Altogether, these data suggested that HOXDeRNA bound to glioma-specific genes and promoted their transcription by titrating out the PRC2 silencing complex.

### Astrocyte transformation leads to the activation of key glioma stem cell super-enhancers

We assembled a list of 174 common GSC super-enhancers (SE) by cross-intersecting three datasets [16] and examined H3K27Ac coverage as a common marker of active SEs in the corresponding genomic intervals in control and HOXDeRNA-transformed astrocytes. We found that almost all GSC SEs were highly enriched with H3K27Ac after, but not before HOXDeRNA-induced astrocyte transformation (Figure 3A). Representative glioma SEs are associated with critical protein-coding and non-coding oncogenes such as SOX2, WEE1, EGFR and miR-21 (Figure 3B). SEs can be activated by TF binding (reviewed by [17, 18]; we, thus, searched for the TF motifs in the H3K27Ac-covered regions of the activated SEs. The binding motifs for OLIG2 and SOX2, two TF marks of GBM subtypes induced by HOXDeRNA and upregulated after transformation, were enriched most highly (Figure 3C), suggesting their regulation of numerous SEs. A comprehensive analysis of SOX2 and OLIG2 ChIP-seq datasets from GSCs revealed that 82% of the SEs were occupied by SOX2, OLIG2, or both (Figure 3D). Of note, ChIRP data analysis indicated that none of the SEs was bound by HOXDeRNA, suggesting that their activation is downstream of the TF derepression.

**Figure 3.**
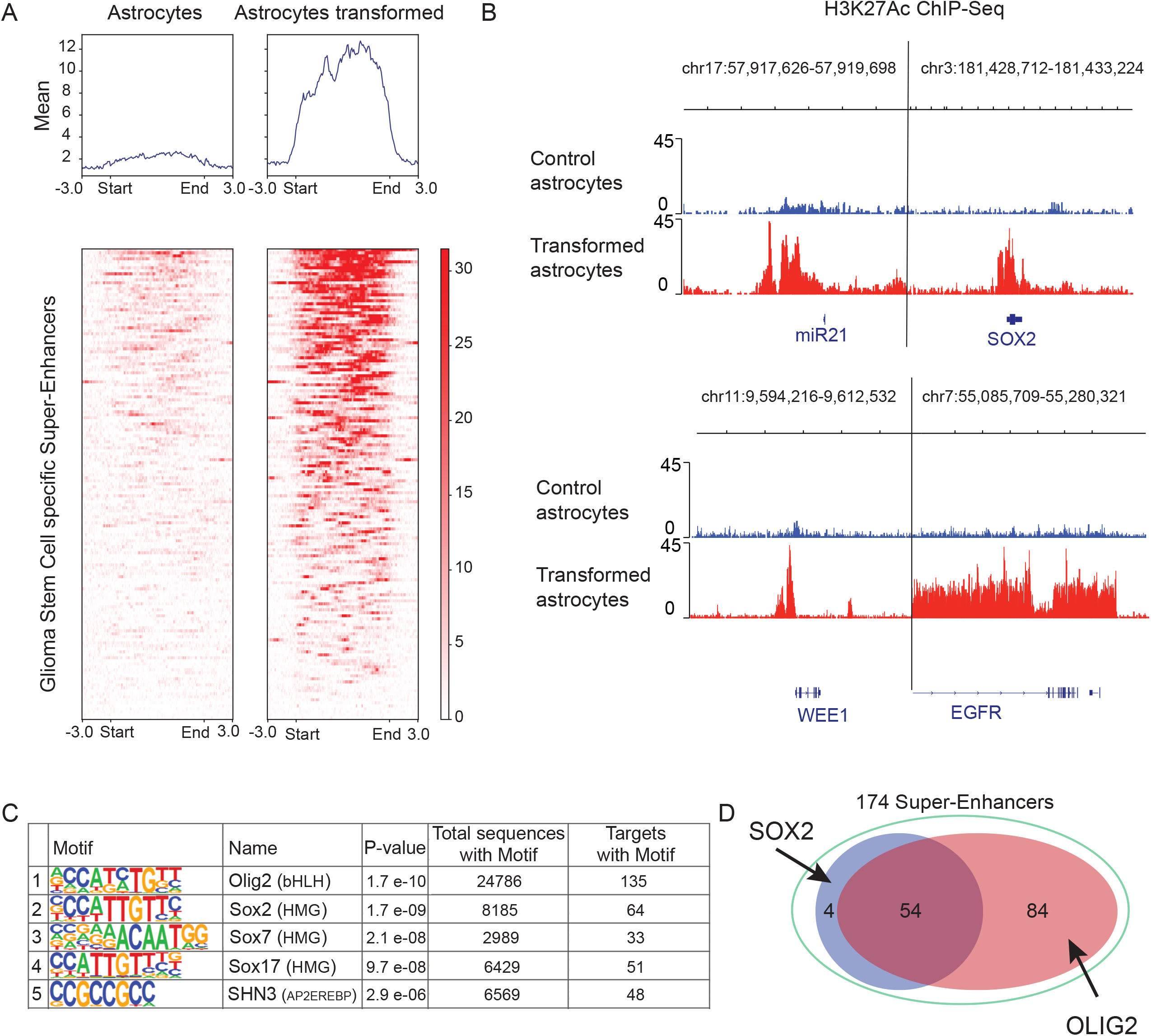
Induction of HOXDeRNA activates GSC-specific super-enhancers. A. Raw H3K27Ac coverage was monitored in control and HOXDeRNA-transformed astrocytes at 174 glioma-specific SEs. For length normalization, the SEs were split into the same number of bins. The average read coverage value per bin across all SEs or the individual value per bin were plotted as line graphs (top) or heatmaps (bottom), correspondingly. The “Start” and “End” marks define the ends of the enhancer. B. H3K27Ac ChIP-seq signals are shown for representative GSC SEs in control and transformed astrocytes. C. The list of the top 5 TF binding motifs enriched in the SE-associated H3K27Ac peaks. TF enrichment analysis was performed with Homer software. D. ChIPseq demonstrates that SOX2, OLIG2, or both bind to 82% of GBM SEs. The lists of SEs bound by SOX2 or OLIG2 were intersected and visualized as Venn diagram.

To further investigate the effects of SE activation on the coordinated transcriptional programs in glioma, we integrated our RNA-Seq and H3K27Ac datasets with the CTCF ChIPseq and HiC datasets (see the legend for Supplemental Figure 4 for the list of tracks and accession numbers). We hypothesized that SEs might concordantly activate clustered genes located within the same topologically associated domains (TADs). We searched for such clusters exhibiting concordant transcriptional activation after astrocyte transformation and located within the same TADs in glioma. We found that HOXDeRNA transformation-induced protocadherin (PCDH) genes were characterized by genomic interactions with glioma-specific SEs. Multiple PCDHB genes located in the same glioma TAD were activated after transformation by the HOXDeRNA-activated SE (Supplemental Figure 4, dotted rectangles mark specific HiC interactions enriched in GSCs compared to astrocytes, SEs are depicted in green). Moreover, these interactions appeared dependent on a cluster of CTCF sites bound by CTCF proteins in glioma cells but not in astrocytes (CTCF tracks, regions highlighted in yellow).

### HOXDeRNA is globally recruited to chromatin by EZH2

Because genome-wide binding of HOXDeRNA was associated with the removal of PRC2 from essential glioma genes upon astrocyte transformation, we tested whether HOXDeRNA bound to EZH2 directly using the ChIRP-WB technique. Our results showed a direct binding between HOXDeRNA and EZH2 (Figure 4A). In parallel, Crosslinking and Immunoprecipitation (CLIP) experiments demonstrated that EZH2 was highly enriched for HOXDeRNA (Figure 4B). We further hypothesized that EZH2 can directly recruit HOXDeRNA to chromatin. We identified 1453 EZH2-covered and 166 EZH2-free genomic regions in astrocytes that gain binding of HOXDeRNA after transformation. We then knocked-down EZH2 by siRNA (Figure 4C) and examined HOXDeRNA binding in these regions using ChIRP-Seq. EZH2 KD strongly reduced HOXDeRNA binding at EZH2-covered genomic regions, but did not affect HOXDeRNA binding at EZH2-free regions (Figure 4D). For example, the binding of HOXDeRNA to the TSS of glioma master TFs (e.g., OLIG2, SOX2, POU3F2, and ASCL1) in GSCs and transformed astrocytes disappeared following the EZH2 KD (Figure 4E). This data indicates that EZH2 is critical for the recruitment of HOXDeRNA to most of its chromatin binding sites.

**Figure 4.**
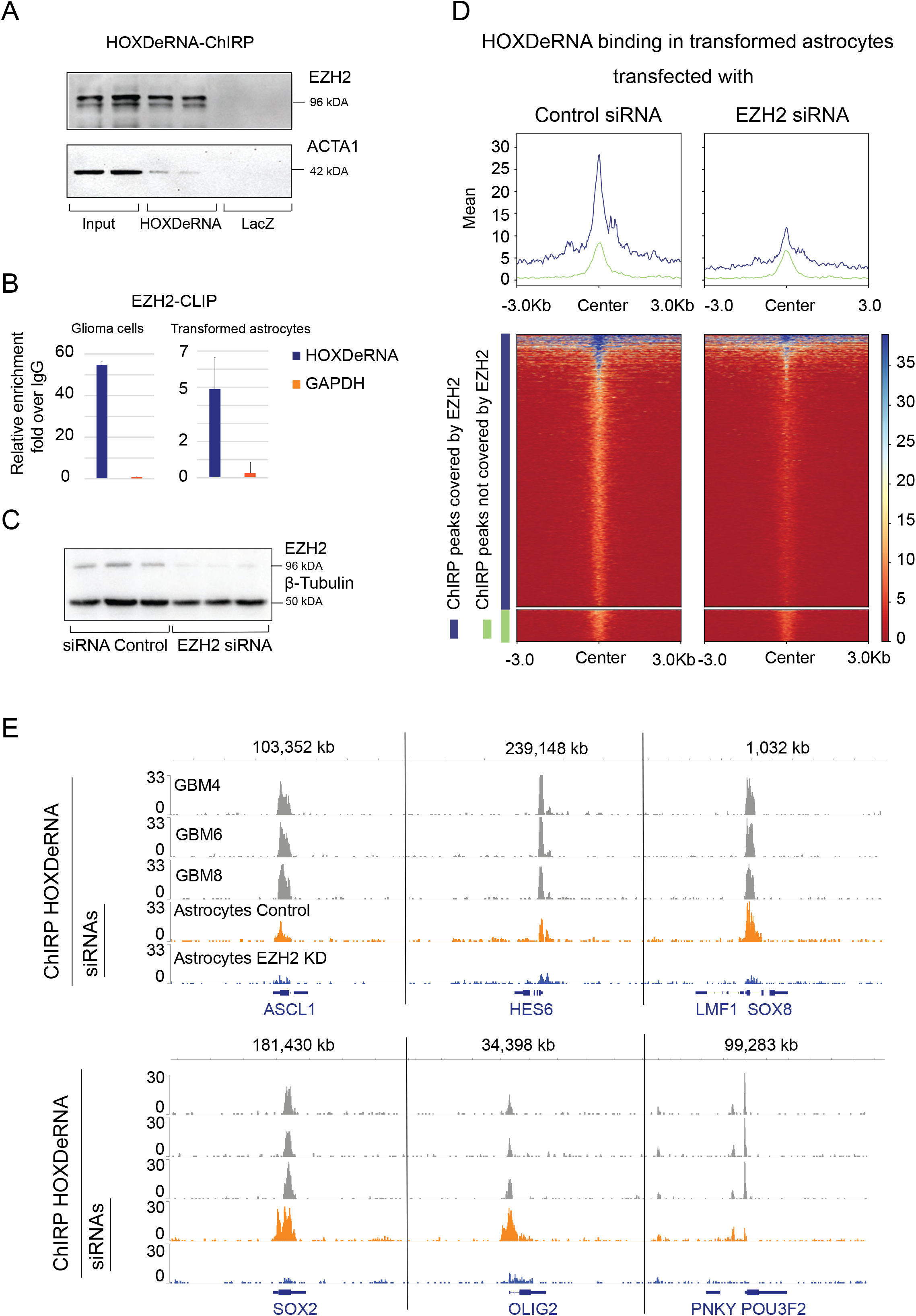
HOXDeRNA genome-wide binding depends on EZH2. A. ChIRP with probes for HOXDeRNA and LacZ (negative control) followed by the Western blots with antibodies recognizing EZH2 and ACTA1 was performed on transformed astrocytes and visualized, with 1% input. Two representative biological replicates per group are shown. B. CLIP with EZH2 and IgG antibodies followed by qRT-PCR detection of HOXDeRNA and GAPDH mRNA was performed in glioma LN229 cells and transformed astrocytes. EZH2/IgG ratios are demonstrated (n=3, mean+ SD). C. Western Blot validating EZH2 inhibition in transformed astrocytes at 48 hours post-transfection with EZH2 siRNAs (n=3). D. ChIRP HOXDeRNA signals were measured in transformed astrocytes transfected with either control or EZH2 siRNAs and visualized as average (line graph, top) or individual values (heatmap, bottom) at HOXDeRNA peaks (center +/- 3 kb). HOXDeRNA binding was analysed separately for the peaks covered or not by EZH2 in control astrocytes (blue and green lines, correspondingly). E. HOXDeRNA binding at the promoters of key glioma master TF genes is shown for GSCs (top three tracks) and transformed astrocytes transfected with either control siRNAs or EZH2 siRNAs (two bottom tracks).

### Binding of HOXDeRNA to EZH2 depends on the RNA quadruplex structure

To investigate if EZH2 interacted with HOXDeRNA via an RNA quadruplex (rG4) structure, as reported for its interaction with other RNAs [9, 10], we first analyzed the capacity of HOXDeRNA to form rG4s structures. The analysis using the QGRS mapper tool (https://bioinformatics.ramapo.edu/QGRS/index.php) yielded 5 putative rG4-forming sequences (Supplemental Figure 5A). We then employed circular dichroism (CD) spectroscopy to assess the ability of these predicted G-rich HOXDeRNA motifs to fold into rG4 structures. An increased CD peak of the RNA oligonucleotides at 263 nm in the rG4-favoring K^+^ environment and a reduction in the corresponding peak intensity in the rG4-unfavoring Li^+^ environment suggests the formation of an rG4 [19]. Among the tested sequences, three oligonucleotides (rG4-1, rG4-2, and rG4-4) showed a typical rG4 CD behavior, suggesting the highly probable formation of the rG4 structures in the cells (Supplemental Figure 5B). To further check the stability of these rG4 structures, we used CD melting approach [20], which identified rG4-1 as the most stable structure with a Tm of 57.5°C (Supplemental Figure 5C). We then tested the ability of potential rG4 sequences to bind to EZH2 and found that rG4-1 exhibited the highest affinity for EZH2 (Figure 5A). These data suggested that EZH2 binding to HOXDeRNA was mediated by the rG4-1 structure.

**Figure 5.**
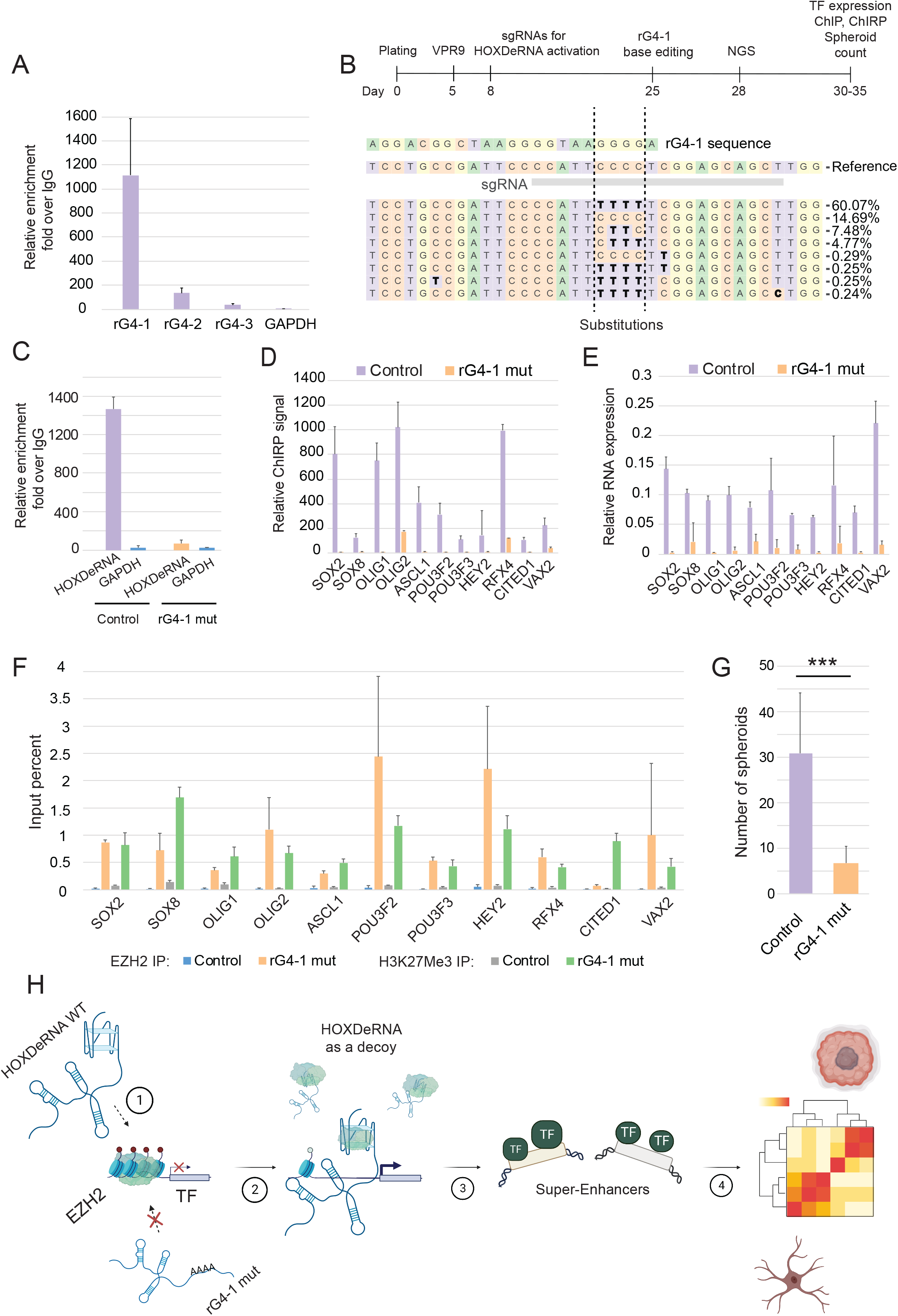
An RNA quadruplex rG4-1 element in HOXDeRNA mediates its EZH2 binding and PRC2 removal, global regulatory activity, and transformation capacity. A. CLIP with EZH2 antibody followed by qRT-PCR detection of three putative HOXDeRNA rG4-containing regions, using GAPDH mRNA as a negative control (mean + SD, n=3). B. Schematic timeline for rG4-1 base editing experiments (top). DNA analysis confirming efficient C-to-T editing in the HOXDeRNA rG4-1 genomic region, corresponding to the G-to-A editing in the HOXDeRNA. Alleles with a substitution rate of > 0.1% are visualized. Substituted nucleotides are shown in bold. C. CLIP analysis of WT and rG4-1-edited cells with EZH2 antibody followed by qRT-PCR for HOXDeRNA and GAPDH mRNA (mean + SD, n=3). D. rG4-1 base editing abolishes the binding of HOXDeRNA to its targets in transformed astrocytes. ChIRP-qPCR analysis of HOXDeRNA/gene promoter binding was performed in control (n=3) and base edited transformed astrocytes (n=3). The data were normalized to the GAPDH gene and presented as bars (mean + SD). E. rG4-1 base editing disrupts the derepression of HOXDeRNA target genes after astrocyte transformation. qRT-PCR analysis of the corresponding set of HOXDeRNA-induced target mRNAs was performed in both control (n=3) and base-edited (n=3) transformed astrocytes and normalized to GAPDH mRNA levels. The data are shown as bars (mean+ SD). F. rG4-1 base editing prevents removal of PRC2 from HOXDeRNA targets after its activation. ChIP-qPCR reactions with EZH2 and H3K27Me3 antibodies on control (n=3) and edited clones (n=3) were normalized for input and shown as bars (mean+ SD). G. rG4-1 base editing inhibits astrocyte transformation. Number of spheroids were quantified in both control (n=3) and base-edited (n=3) transformed astrocytes (mean + SD). H. A model of genome-wide function of HOXDeRNA. rG4-dependent HOXDeRNA binding to EZH2 and recruitment to PRC2-silenced promoters in astrocytes (1) leads to reduced PRC2 repression in the corresponding chromatin regions, gene derepression, and active state of glioma master TFs (2), followed by SE activation (3) and further transcriptional reprograming (4).

To further investigate the role of rG4-1 in the HOXDeRNA function, one of the two G-nucleotide stretches forming the rG4-1 structure was edited using the CRISPR base editing technique [21] (Figure 5B). We tested whether these rG4-1-abolishing G-to-A substitutions within the HOXDeRNA affected its binding to EZH2 and transformative effects on astrocytes. Indeed, rG4-1 ablation reduced HOXDeRNA binding to EZH2 about 20 fold (Figure 5C) without affecting HOXDeRNA expression (Supplemental Figure 5D). As a downstream readout, we selected a panel of marker genes repressed by PRC2 in astrocytes but bound by HOXDeRNA and transcriptionally activated after transformation. The results indicated that rG4-1 ablation abolished HOXDeRNA binding to the gene promoters, strongly reduced corresponding EZH2 and H3K27Me3 marks, and the genes’ expression (Figure 5D-F). In accordance with these molecular alterations, the rG4-1-deficient astrocytes exhibited a reduced transformation capacity as observed by the matrix-independent spheroid growth (Figure 5G).

We propose a model in which HOXDeRNA is recruited to essential glioma genes in rG4-1/EZH2 -dependent manner and keeps them active by operating as a PRC2 decoy (Figure 5H. HOXDeRNA regulates a dynamic balance between the activating and repressive molecular machineries by preventing the accumulation of PRC2 in transformed astrocytes and glioma cells to the levels critical for gene silencing.

## Discussion

There are thousands lncRNAs transcribed from human genome, they are often poorly conserved and cell-type specific, especially enhancer-associated lncRNAs (eRNAs) [22, 23]. Many lncRNAs are associated with chromatin-modifying complexes and may affect gene expression [24, 25]. Nevertheless, the functions of only a few lncRNAs have been demonstrated (reviewed in [26, 27]) and roles of lncRNAs in cell fates and plasticity are largely unknown. Here we report a broad genome-wide function of a single eRNA that binds to and derepresses promoters of genes encoding glioma master TFs scattered throughout the genome. These TFs in turn activate multiple glioma-specific SEs and their associated target genes, including protein-coding genes, miRNAs, and lncRNAs, involved in the malignant phenotype of glioma. Among them are the major glioma factors such as EGFR, PDGFR, TERT, miR-21, miR-10b, and HOXD-AS2. Such a global transcriptional regulatory role of a lncRNA has been so far only demonstrated for a few lncRNAs, which recruit gene-silencing or activating epigenetic factors to multiple genomic loci. Among them are XIST [28–30], LINC-PINT [31], NRIP1e [32], and KCNQ1OT1 [33] [34], and they mostly function *in cis*. To our knowledge, only one lncRNA (lncPRESS1) working on multiple gene targets *in trans* as a decoy for specific chromatin modifiers has been reported thus far [35]. However, molecular determinants of lncPRESS1 binding to a deacetylase SIRT6 and the repertoire of direct gene targets regulated directly by this lncRNA are yet unknown. We describe the first lncRNA that is recruited to its gene targets in an rG4/EZH2 -dependent manner and drives a cancer-specific transcriptional program by removing PRC2 repression from the key elements of glial regulatory circuitry, altering the epigenetic landscape across all 23 chromosome pairs, and resulting in astrocyte transformation to glioma-like cells. Most notably, the expression of almost all glioma master TFs appears under direct regulation of HOXDeRNA.

Master TFs, SEs, and their gene targets define cell fates and identities in normal and disease states. Overexpression of a few master TFs of pluripotency has been shown to induce significant transcriptomic and phenotypic alterations through the processes of cell reprogramming and transdifferentiation [36, 37]. The list of glioma-specific TFs includes 50 factors [1]. However, a minimal core of only four neurodevelopmental TFs (SOX2, OLIG2, POU3F2, and SALL2) appears sufficient for the reprogramming of differentiated glioma cells into a cell population that recapitulates the epigenetic and transcriptomic programs of patient derived GSCs [15]. This small set of TFs is also sufficient to confer tumour-propagating properties *in vivo*. Remarkably, our data indicate that activation of HOXDeRNA in astrocytes leads to its direct binding to promoters of 44 out of 50 glioma-specific TFs, all silenced in astrocytes by PRC2, and derepressing their expression. Among the HOXDeRNA-activated TFs are the four core TFs and 12 additional TFs enriched in stem-like tumor-propagating cells relative to both astrocytes and differentiated glioma cells (OLIG1, SOX8, ASCL1, HES6, POU3F3, HEY2, SOX5, RFX4, KLF15, CITED1, VAX2, MYCL1) [15]. Of note, there are two distinct GSC groups driven by key TFs (i.e., OLIG1/2 and RUNX1/2/TFAP2A) that define proneural and mesenchymal GBM subgroups, respectively [38, 39]. Our data indicate that both groups of the TFs are directly activated by HOXDeRNA, suggesting its involvement in both proneural and mesenchymal transcriptional programs. Furthermore, OLIG2 and SOX2 binding motifs are found in 82 % of HOXDeRNA-activated SEs, many of which serve as enhancers of glioma-driving genes (e.g., OLIG2, SOX2, EGFR, BRD4, POU3F2, miR-21, and others). This data supports the idea of the reciprocal relationship between TFs and SEs, where TF can be both a regulator and a target of the SE. Such scenario has been observed, for example, in embryonic stem cells, pro-B cells, myotubes, Th cells and macrophages [40–42]. Most importantly, the data indicate that a compact glioma-specific auto-regulatory transcriptional network, anchored on TFs and SEs, is epigenetically controlled by an oncogenic eRNA that serves as a PRC2 modulator.

The components of PRC2 repressive complex, including EZH2, bind RNA *in vitro* and *in vivo* [4, 9]. This versatile promiscuous binding can be rationalized by the fact that PRC2 recognizes short G-tracts and G-quadruplexes, small motifs that are ubiquitous across the transcriptome [3]. This has led to the speculation that G-rich single-stranded or secondary RNA structures, common to multiple RNAs, may be involved in RNA-PRC2 interactions. Recent PRC2 CLIP studies have shown that 8-mer G-tract sequences are also enriched at cross-linking sites *in vivo*. Moreover, tethering G-tract or G-rich RNAs to the 5’ end of genes removes PRC2 components and H3K27Me3 from chromatin [10]. Consistent with these reports, we demonstrate that HOXDeRNA binds to EZH2, and their interaction is mediated by the rG4 structure. By using CRISPR base-editing, which does not affect the activation and transcription of HOXDeRNA, we for the first time provide the evidence of rG4-EZH2 binding and its global impact on gene expression in the *in vivo* chromatin context preserved by chemical cross-linking. According to our model, HOXDeRNA removes PRC2 in an rG4-dependent manner from essential glioma-driving genes during the process of astrocyte transformation and prevents the accumulation of PRC2 on these genes in glioma, thereby regulating the dynamic balance between PRC2 and transcriptional machineries. Our analyses were performed on the bulk chromatin material, and a more detailed single-cell-based approach would be required to refine this model. Furthermore, HOXDeRNA binding to EZH2 has also been reported in other cancers, such as colorectal and osteosarcoma [43, 44]; thus, it would be important to investigate its genome-wide regulatory function in non-glioma contexts more broadly. Overall, our results contribute to a growing body of knowledge regarding the functional interplay between PRC2 and RNA and underscore the importance of further investigating the mechanisms underlying these interactions in the context of gene regulation and cellular function.

GBM is a highly heterogeneous disease characterized by a diverse array of mutations. The most frequently observed mutations occur in tumour suppressor genes such as NF1, CDKN2A, PTEN, RB1, and TP53, as well as in genes involved in the regulation of telomere length maintenance (TERT, ATRX, DAXX) and metabolism (IDH1/2) [45, 46]. Recently, research has increasingly focused on the role of epigenetic alterations, including DNA methylation, histone modifications, and chromatin topology in the pathogenesis of this cancer. These alterations impact gene expression through the interplay of cis-regulatory elements, such as promoters, enhancers, and silencers, with trans-acting factors, such as TFs. Since mutations in the HOXDeRNA/HOXD region are very rare and are not associated with the HOXDeRNA activation [11], our study indicates that transcriptional reprogramming underlying neoplastic transformation can occur without genetic alterations. Moreover, multiple cancer drivers beyond glioma TFs, including TERT, EGFR, BRAF, and PDGFR that are established therapeutic targets, are simultaneously upregulated by HOXDeRNA. Many of them are directly derepressed by the HOXDeRNA binding. This observation extends our current understanding of gliomagenesis and glioma biology, which has traditionally been viewed through the lens of mutational landscapes.

The development of novel therapeutic strategies for GBM is a pressing issue, as the current standard care fails to improve patients’ survival post-diagnosis beyond 15-20 months. The ongoing clinical trials targeting, for example, EGFR, TGF-β, and VEGF-A face conceptual challenges. Among them is the expression of these targets in healthy brain tissues that may be associated with severe brain toxicity of the targeted therapies. Additionally, the heterogeneous nature of GBM cells and the wide expression and activity of these targets may lead to the selection of the antigen-negative or therapy-resistant tumour cells. Our research offers a new approach to tackle these issues by identifying a unique molecular target that is absent in normal neuroglial cells and neuroprogenitors of the brain. This target, a powerful eRNA acting both *in cis* and *in trans*, globally reorganizes chromatin, activates developmental programs, promotes the identity of glioma cells and, thereby, drives gliomagenesis through a transcriptional axis consisting of transcription factors and super-enhancers. As a result, it controls multiple “druggable” (e.g., EGFR, PDGFR, BRAF) as well as “poorly-druggable” (e.g., TFs responsible for GSC tumor-initiating and therapy-resistance properties) factors essential for glioma viability and recurrence. Correspondingly, HOXDeRNA KD is detrimental for glioma growth [11, 47–51]. Therefore, development of HOXDeRNA-targeting therapeutic strategies can lead to bench-to-bed translation and provide an important avenue complementing fundamental molecular studies.

## Methods

### Resource Availability

#### Lead Contact

Further information and requests for resources and reagents should be directed to the Lead Contact Dr. Anna M. Krichevsky, Brigham and Women’s Hospital and Harvard Medical School, 60 Fenwood Road, Boston, MA 02115, USA.

#### Materials Availability

This study has not generated new unique reagents.

#### Data and code availability

Sequencing data generated in this study have been deposited to GEO with accession number GSE227805.

### Cell cultures and transfections

Human cells were used in accordance with institutional review board guidelines at Brigham and Women’s Hospital. Low-passage human GBM stem cells (GBM4, GBM6, and GBM8) were a generous gift from Dr Hiroaki Wakimoto, MGH. The tumorigenic, genetic, and molecular properties of these cells have been described previously [53]. Cells were maintained in serum-free neurobasal media supplemented with N-2 and B-27 Plus Supplements (Gibco™), 3 mM Gibco® GlutaMAX™ Supplement, 50 units/ml penicillin and 50 units/ml streptomycin (Gibco™), 2 μg/ml heparin (Sigma-Aldrich), 20 ng/ml FGF2 (Sigma-Aldrich), and 20 ng/ml EGF (Sigma-Aldrich). Cells were passaged by dissociation using the Neurocult Stem Cells chemical dissociation kit (Stem Cells Technologies). Normal human astrocytes immortalized by E6/E7/hTERT (a generous gift from Dr. Yukihiko Sonoda [12] were maintained in Neurobasal medium similarly to GSCs. Human primary astrocytes were cultured as previously described [11]. Cell transfections with EZH2 siRNAs (50nM) have been performed as previously described [11]. Cell cultures were regularly tested for mycoplasma and cell lines were authenticated.

### CRISPR activation of HOXDeRNA

Lentiviral plasmids to generate stable dCas9-VPR nuclease-expressing cell populations, Edit-R CRISPRa lentiviral sgRNA non-targeting control, and two custom-made Edit-R CRISPRa human HOXDeRNA lentiviral sgRNAs, all from Dharmacon, were employed as previously described [11]. sgRNAs used for HOXDeRNA gene activation were: AAGGCGCAGGCTGGCAGTTC, CCAGCCTGCGCCTTTGCAGC. Edit-R CRISPRa lentiviral sgRNA non-targeting control (cat. GSGC11913) was used as control sgRNA. NHA cells were sequentially transduced with dCas9-VPR followed by sgRNAs according to the Dharmacon technical manual.

### Chromatin Immunoprecipitation followed by DNA Sequencing (ChIP-Seq) and data analysis

ChIP-Seq was performed using the SimpleChIP® Enzymatic Chromatin IP Kit (Magnetic Beads) #9003. Briefly, 10 million cells were cross-linked with 1% formaldehyde and washed twice with ice-cold PBS. The collected pellet was resuspended in 2 ml RIPA buffer (BP - 115X, Boston BioProducts) with protease inhibitors and fragmented to ∼300 bp using MISONIX S-4000 Sonicator (amplitude 30%, 30 sec ON / 30 sec OFF, 30 min). 20 μg of chromatin in 1 ml of IP dilution buffer (16.7 mM Tris-HCl pH 8, 0.01% SDS, 1% Triton X-100, 167 mM NaCl, 1.2 mM EDTA) per ChIP was mixed with 10 μg of the following antibodies: H3K27Ac (#4353), H3K27Me3 (#9733) (all from Cell Signaling), EZH2 (Cat. 39002, Active Motif). The IP mixes were incubated overnight under rotation at 4 °C, followed by an additional 4 hours with Dynabeads™ Protein G (30 μl per sample). IPs were washed twice in a low salt buffer (10 mM Tris-HCl pH 8.0, 1 mM EDTA pH 8.0, 150 mM NaCl, 1% Triton X-100 in distilled water), once with a high salt buffer (10 mM Tris-HCl pH 8.0, 1 mM EDTA pH 8.0, 500 mM NaCl, 1% Triton X-100 in distilled water) and once in TE buffer, eluted and de-crosslinked according to the instructions (SimpleChIP® Enzymatic Chromatin IP Kit). DNA was purified using the Monarch PCR and DNA Clean Up Kit (#T1030L, NEB) and ChIP-seq libraries were prepared using the NEBNext® Ultra™ II DNA Library Prep Kit for Illumina® (E7546S, NEB) and sequenced on the Illumina HiSeq 2500 platform configured for 50-bp single-end reads.

For data analysis, Fastqsanger files were aligned to the human reference genome (GRCh37/hg19) using Bowtie2 with default parameters. PCR duplicates were removed using samtools rmdup version 1.13 [54]. The resulting aligned Bam files were transformed to Bigwig format without scaling or normalization using deeptools version 3.5.0 [55]. Bed files for peaks were created using MACS2 (version 2.2.7.1, [56]. The IGV web application [57] was used for visualization. The GSC specific list of SEs was generated as a bed file by intersecting three glioma SE bed files from GSE121601. This list intersected with our H3K27Ac data was used to produce Figure 3A.

For Figure 3C, SOX2 (GSM1306360_MGG8TPC.SOX2, n=2) and OLIG2 (GSM1306365_MGG8TPC.OLIG2, n=2) BED files (peaks) were downloaded from GEO. All overlapping intervals were merged with MergeBED function (BEDtools,[58]) after replicates were concatenated tail-to-head. “Intersect intervals” function (BEDtools) was used to generate lists of SOX2-bound and OLIG2-bound super enhancers. Resulting lists were intersected and visualized as Venn diagram.

ChIP-qPCR data was analysed in 4 steps: 1) 1% input Ct values were adjusted to 100% by subtracting 6.64; 2) Ct (IP) was subtracted from the adjusted input Ct to obtain delta Ct; 3) calculate 100*2^delta CT for each IP; 4) calculate a mean with SD for each condition.

### Chromatin Isolation by RNA purification followed by DNA Sequencing (ChIRP-Seq) and data analysis

ChIRP was performed based on the previously published protocol [59]. Briefly, 48 biotinylated oligonucleotide probes specific for nascent HOXDeRNA RNA were designed and manufactured using ChIRP probe designer from LGC Biosearch Technologies (https://www.biosearchtech.com/support/tools/design-software/stellaris-probe-designer).

Probes for LacZ mRNA (Sigma-Aldrich, 03-307) were used as negative control. Single cell suspensions were prepared from 100 million adherent or suspension cells followed by cross-linking with 2% glutaraldehyde (Sigma, G5882-50), and centrifugation at 20 rpm RT for 20 minutes. The reaction was quenched with glycine solution (7005S, Cell Signaling), and the cells were washed twice with ice-cold PBS. Sonication was performed in 15 ml falcon tubes with RIPA buffer and protease inhibitors (cOmplete Mini EDTA free), and chromatin was fragmented to ∼200 bp using MISONIX S-4000 sonicator (amplitude 30%, 30 sec ON / 30 sec OFF, 20 min). The chromatin from each cell line was mixed with the protease and RNAse inhibitors (SuperaseIn RNAse Inhibitor, AM2696, ThermoFisher Scientific), split into three tubes (for input, LacZ, and HOXDeRNA binding), incubated for 4 h at 37C 1000 rpm with respective sets of biotinylated probes, and followed by another 2 h incubation with Dynabeads MyOne Streptavidine C1 (62001, ThermoFisher). The beads were washed twice with low and high salt buffers at 37C, 1000 rpm, followed by de-crosslinking with Proteinase K for 4 hours at 65C. For ChIRP-WB analysis, the beads were loaded into NuPAGE Bis-Tris protein gels (Invitrogen, NP0335PK2). EZH2 (Cat. 39002, Active Motif, dilution 1:1000) and ACTA1 (Catalog # A00885-40, GenScript, dilution 1:500) antibodies were used for protein detection. For sequencing, DNA was purified using the Monarch PCR and DNA Clean Up Kit (#T1030L, NEB), Illumina NGS libraries were prepared using the NEBNext Ultra II DNA Kit (NEB, E7645S), and sequenced for 50-nt SE reads. For data analysis, Fastqsanger files were mapped to hg19 genome using bowtie2 (default settings), and the resulting aligned Bam files were transformed to Bigwig format using deeptools version 3.5.0 [55]. Raw reads were quantified for each 50-nucleotide bin for each Bigwig file. The Bigwigs signal tracks were visualized with IGV browser. Peaks were called using MACS version 2.0.10 [56] (default parameters). The resulting bed files were intersected with other interval data. For Figure 2A, ChIRP peaks were annotated (using Homo_sapiens.GRCh37.75.gtf (hg19)) with ChIPseeker (version 1.34.1, [60]) to the nearest genomic feature. The number of peaks corresponding to various genomic features was plotted as a Venn diagram. For Figure 2B, ChIRP peaks were annotated to the nearest gene for each cell line using ChIPseeker, followed by visualization of intersected gene lists with a web tool ’Intervene’ [61]. Average profile plots and heatmaps for Figures 2C-F, 3A; 4C, and Supplemental Figure 2 were generated using the deeptools functions plotProfile and plotHeatmap [55].

The ChIRP-qPCR data were analyzed using 2^-delta Ct formula, where delta Ct= Ct (IP) - Ct(GAPDH). Primers used in the reactions are listed in Supplemental Table 1.

### mRNA-Seq, qRT-PCR, and data analysis

Total RNA was isolated with Norgen Biotek kit followed by treatment with DNAse I (Promega) at 37°C for 30 minutes. qRT-PCR reactions were performed as previously described [11], with primers listed in the Supplemental Table 2. The data were analysed in three steps: 1) Ct (gene of interest) was subtracted from GAPDH Ct (normalization control) to obtain delta Ct; 2) 2^-delta Ct was calculated for each sample; 3) means and SD for each group were calculated and plotted as bars. Alternatively, cDNA libraries were prepared, and deep sequencing was performed by Novogene. For the analysis of RNAseq data, Fastqsanger files were trimmed to remove Illumina adapters with Trimmomatic (version 0.39, [62], and aligned to the GRCh37/hg19 genome using HISAT 2.2.1 [63] with the annotation Homo_sapiens.GRCh37.75.gtf (hg19) containing exon-exon splice junctions. Raw counts were measured with featureCounts (part of the subread 2.0.0 package, [64]). Differential expression analysis was performed using Deseq2 (version 1.39.4, [65]. PCA plot and GO analysis for Supplemental Figure 1A were produced and visualized using Debrowser (version 1.24.1, [66]. Heatmaps for Figure 1B and Supplemental Figure 3 were generated with heatmap.2 function of the R gplots package, with z-scores calculated for each gene across all samples (version: 3.1.3, [67]).

Principal Component Analysis (PCA) based on machine learning selected features for Figure 1D was performed as following. We used the TCGA TARGET GTEx dataset from the UCSC RNA-seq compendium, where TCGA, TARGET, and GTEx samples were processed using the same bioinformatic pipeline, including alignment to the hg38 genome, and gene expression calling with RSEM [68] and Kallisto [69] methods. Uniform processing eliminated computational batch effects in this dataset. The dataset can be downloaded at: https://xenabrowser.net/datapages/?cohort=TCGA%20TARGET%20GTEx&removeHub=https%3A%2F%2Fxena.treehouse.gi.ucsc.edu%3A443. To make our RNA-Seq data directly comparable to the selected datasets, they were processed through the same pipeline.

Feature selection and training for generating Fig. 1D was performed as following. To define DEG between the groups, we used a significance threshold of p < 0.05, followed by a selection of the top first 500 genes sorted by fold change. Feature selection was then applied to these 500 genes using the SHAP package [70]. We calculated the importance of the genes using the Shapley values and used this information to improve the performance of the model by removing the less important genes. In the final step, we selected 30 genes with the highest scores for model prediction. To account for possible batch effects of different datasets, we transformed the data for the selected genes into a rank order before training the model. We used an xgboost model [71] to predict whether the samples were associated with normal brain categories (Forebrain (n=857), Cerebellum (n=214), Midbrain (n=57)), LGG (n=522), or GBM categories (Mesenchymal (n=54), Classical (n=41), Proneural (n=39), and Neural (n=28)).

### CRISPR base editing: plasmids, cell transfection, and data analysis

pCMV_BE4max_P2A_GFP plasmid was a gift from David Liu (Addgene plasmid # 112094; http://n2t.net/addgene:112094; RRID: Addgene_112094). Plasmids encoding sgRNA for HOXDeRNA rG4-1 base editing (5’CCATTCCCCTCGGAGCAGCT) and control sgRNA were purchased from Vectorbuilder (pLV[gRNA]-EGFP:T2A:Puro-U6). Six-well plates of transformed astrocytes in Neurobasal medium were transfected with 6 ul Lipofectamine 2000, 750 ng pCMV_BE4max_P2A_GFP, and 250 ng sgRNA per well. Cell culture medium was changed 4 hours later, and Puromycin (100 µg/mL) was used for cell selection 24 hours post-transfection. Genomic DNA was extracted using the genomic DNA purification kit (Dneasy Blood and Tissue kit, Cat. 69504, Qiagen), the specific fragment containing rG4-1 sequence was amplified using Phire Green Hot Start II PCR Master Mix (F126S) with the forward primer 5’TCCGCCTGGAAAAGAAGTCC and reverse primer 5’GAGGCAAGACTTTGGTGGGA, and the purified amplicon was sequenced at MGH DNA core facility.

### Cross-linking and immunoprecipitation (CLIP)

CLIP experiments were performed based on previously published protocol [72]. Briefly, to covalently cross-link proteins to nucleic acids, 2 × 10^7^ cells were exposed to UV irradiation (200 mJ/cm²) for 2 minutes. The cells were then lysed with RIPA buffer containing Protease Inhibition Cocktail and RNase Inhibitor. Immunoglobin-coated magnetic protein beads (Thermofisher Scientific) were incubated with EZH2 (Cat. 39002, Active Motif) or IgG (Catalog # 2729S, Cell Signaling) antibodies. The complexes were washed with RIPA buffer and the samples were treated with DNAse I (Promega) for 30 minutes at 37°C, followed by Proteinase K (10% SDS and 10 mg/ ml Proteinase K in RIPA buffer) treatment for 30 minutes at 37°C with shaking. RNA was further isolated with a phenol:chloroform:isoamyl alcohol (25:24:1) solution, precipitated with isopropanol, and resuspended in RNase-free water. qRT-PCR was performed for nascent HOXDeRNA and GAPDH mRNA. Primers are listed in Supplemental Table 2. Fold enrichment over IgG was calculated in two steps: 1) DDCt = (Ct IP) - (Ct IgG), and 2) fold enrichment = 2^-DDCt.

### Circular dichroism (CD)

The oligonucleotides were dissolved in 150 mM K^+^ in T_10_E_0.1_ buffer (10 mM Tris-HCl, 0.1 mM EDTA). 200 µL of 10 µM solution was placed in quartz cuvettes (1 mm path length) and the spectra were collected in the range between 200 and 320 nm at 20 °C from three scans, and a buffer baseline was subtracted from each spectrum. Increased peak intensity of the oligo under the K^+^ environment (compared to the Li^+^ environment) at 260-265 nm and a trough at around 240 nm suggests the formation of an rG4. JASCO J815 spectropolarimeter was used to collect CD spectra. The data was plotted using GraphPad Prism (Smooth: 10 neighbors in each side, second order smoothing polynomial).

### CD melting

CD melting experiments were performed in the same spectrophotometer and similar solution conditions as described above. The CD ellipticity of 10 µM folded solution of rG4s was tracked at 263 nm using variable temperature measurement method. The thermal data was collected between 20 °C to 95 °C sampling per °C at a rate of 1°C/ min. The data was plotted in a XY format using GraphPad Prism (Smooth: 6 neighbors in each side, fourth order smoothing polynomial) and first derivative of the data was plotted to calculate the melting temperature (T_m_) of the structure, which is defined as a temperature at which the sequence is equally populated in the folded and unfolded states.

### Quantification and Statistical analysis

All statistical analyses were performed using Prism 8 GraphPad Software. Statistical significance of the differences between groups was measured with two-tailed unpaired t-test. In figure legends: *p< 0.05, **p<0.01, ***P<0.001, ****p<0.0001. Non-significant differences are marked with NS.

**Additional material 1.** Supplemental Figures 1-5.

**Additional material 2.** Supplemental Table 1. Differentially expressed genes for Figure 1B.

**Additional material 3.** Supplemental Table 2. Primers for mRNA and HOXDeRNA detection by qRT-PCR or Clip-qPCR. Related to Figures 4B, 5A, 5C and 5E.

**Additional material 4.** Supplemental Table 3. ChIRP-qPCR primers. Related to Figure 5C.

**Supplemental Figure 1. HOXDeRNA is not expressed in normal neuroglial cells, in contrast to glioma cells, and its activation globally alters transcriptional programs. Related to Figure 1**.

A-B. HOXDeRNA expression monitored by RNA-Seq in normal brain cells (GSE122701, GSE157461, GSE166847, GSE73721 and GSE119834 datasets) (A) and GBM tissues, GSCs (GSE119834), and glioma Cancer Cell Line Encyclopedia [52] (B). HOXDeRNA transcript, composed of 3 exons, is expressed in all glioma samples but not in the normal cells of the brain.

A. Transcriptomic differences among primary astrocytes (n=3) and immortalized astrocytes naive or transformed by HOXDeRNA (n=3), visualized as PCA plot.

**Supplemental Figure 2. Genome-wide binding of HOXDeRNA is associated with exclusive PRC2 removal from the transformation-induced genes. Related to Figure 2**.

A. HOXDeRNA binds to all chromosomes. Relative chromosome length and number of HOXDeRNA peaks were calculated for each chromosome as percent of total.

B, C. ChIRP-seq raw read coverage signal, representing the HOXDeRNA binding at the TSS (+/- 5Kb) of the reverse strand of genes downregulated or upregulated after astrocyte transformation, is visualized as an average value for each group (B) or for each individual gene (C). The heatmap is accompanied by a colour scheme indicating the value of the raw read counts.

D, E. Epigenetic status of genes upregulated or downregulated in transformed astrocytes. H3K27Ac, H3K27Me3, and EZH2 ChIP-seq raw signals covering gene bodies of the negative strand (+/- 5Kb) were normalized to gene length and counted, followed by visualization of the average signal for 3 groups: genes upregulated after astrocytes transformation and bound by HOXDeRNA, downregulated after astrocyte transformation and HOXDeRNA-free, unchanged genes (D). Read coverage values for every individual gene body were visualized as heatmaps for negative strand genes, which are enriched in control or transformed astrocytes (E).

F, G. Genes encoding glioma master TFs are bound by HOXDeRNA after astrocyte transformation. HOXDeRNA ChIRP-Seq raw read signal was normalized to gene length and visualized as mean value (F) or individual values (G) for each TF gene body (+/- 10Kb) on forward and reverse strands. The colour scheme represents the values of the raw read counts. H, I. PRC2 is removed from genes encoding glioma master TFs after astrocyte transformation. The EZH2 and H3K27Me3 raw read coverage signal was normalized to gene length and visualized as mean value (H) or individual values (I) for each gene body (+/- 10Kb) on both forward and reverse strands. The colour scheme represents the values of the raw read counts.

J. High-sensitivity H3K27Me3 and EZH2 ChIP-seq tracks in transformed astrocytes visualized for selected glioma master TF genes indicate incomplete removal of PRC2 activity in bulk chromatin after transformation. A different scale is presented in Fig. 2G bottom tracks to visualize relative levels in both non-transformed and transformed astrocytes.

**Supplemental Figure 3. HOXDeRNA binds to and removes PRC2 repression from multiple glioma master TFs. Related to Figure 2**.

HOXDeRNA ChIRP-seq tracks in GSCs and transformed astrocytes, aligned with PRC2 (H3K27Me3, EZH2) ChIP-seq coverage, before and after astrocyte transformation, and visualized for glioma master TF genes.

**Supplemental Figure 4. Integration of astrocytes RNAseq and ChIPseq data with ENCODE and GEO datasets suggests a mechanism of Protocadherin (PCDHB) family upregulation in GBM. Related to Figure 3**.

List of tracks from top to bottom: 1) Protocadherin genes annotated with Refseq. A heatmap visualizing PCDHB2-16 RNAseq-based expression in control and transformed astrocytes, transformed to z-scores; 2-4) SE tracks in three GCSs (GSE121601); 5-8) CTCF ChIP-seq tracks from ENCODE database for GBM (GSM822303) and three types of astrocytes (GSM733765, GSM1022662, GSM749696). Genomic region with differential CTCF binding is highlighted in yellow; 9-10) H3K27Ac coverage before and after astrocyte transformation shows activation of GBM specific SEs (dotted lines); 11-14) 4 HiC maps for GSCs G583, G567, G523 from [16], available for download at https://wangftp.wustl.edu/hubs/johnston_gallo/), and astrocytes (GSE105194). A subset of HiC contacts representing SEs-protocadherins interactions are highlighted with dotted rectangles.

**Supplemental Figure 5. Characterization of rG4 candidates. Related to Figure 5**.

A. Five putative rG4 forming sequences detected with QGRS mapper tool. Search parameters: QGRS max length: 45, min G-Group size: 2, loop size: from 1 to 36.

B. CD spectra of five putative rG4 forming sequences (rG4-1, rG4-2, rG4-3, rG4-4, and rG4-5), along with one non-rG4 sequence. RG4-1, rG4-2, and rG4-4 exhibit common rG4 CD features (a higher intensity peak at 263 nm under K^+^ environment).

C. CD-melting curves indicating the stability of predicted rG4 structures. rG4-1 showed the highest melting temperature of 57.5 °C.

D. Base editing of rG4-1 does not alter HOXDeRNA expression levels in transformed astrocytes. qRT-PCR analysis, normalized to GAPDH mRNA levels (mean + SD, n=3).

## ACKNOWLEDGEMENT

This work was supported by the R21 NS098051, AG060019, R01 CA215072, and R01 NS113929 grants to AMK, and R01GM146997 and R01GM126150 grants to PI. We thank the members of the Krichevsky laboratory and Dr. Leonid Mirny for helpful discussions and valuable insights. This manuscript was edited at Life Science Editors.

## AUTHOR CONTRIBUTION

AMK and ED conceived and designed the study; ED performed most experiments, data analysis, and visualization; PK performed CD and melting curves experiments; AK assisted with computational analysis; PI contributed reagents and advice. AMK supervised the work and acquired funding. ED and AMK wrote the manuscript and all authors revised and approved the manuscript.

