## Supplemental Figures 1-5 for "HOXDeRNA activates a cancerous transcription program and super-enhancers genome-wide"

### Supplemental Figure 1

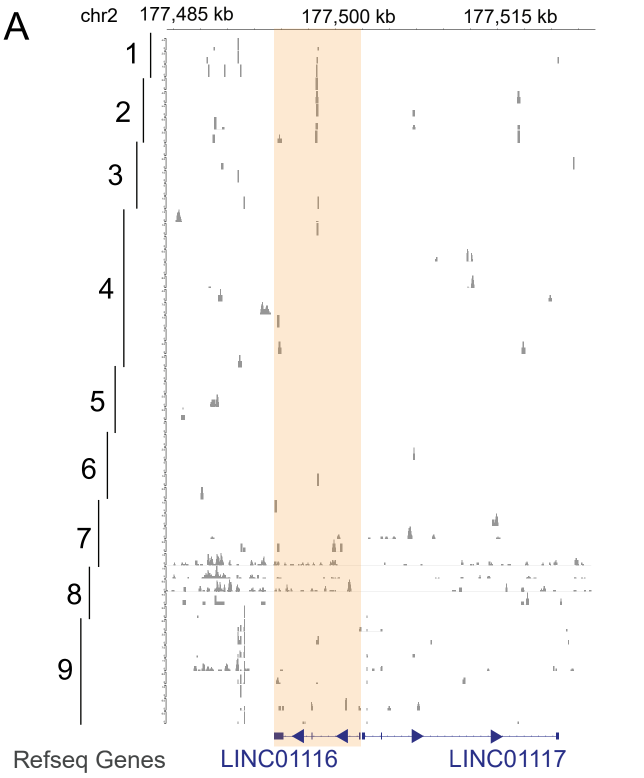


1. Human fetal cortical astrocytes
2. Primary human astrocytes from cerebral cortex
3. Human primary astrocytes
4. Astrocytes from adult temporal lobe
5. Astrocytes from hippocampus
6. Astrocytes from fetal cortex
7. Oligodendrocytes from adult temporal lobe
8. Whole cortex from adult temporal lobe
9. Neural stem cells

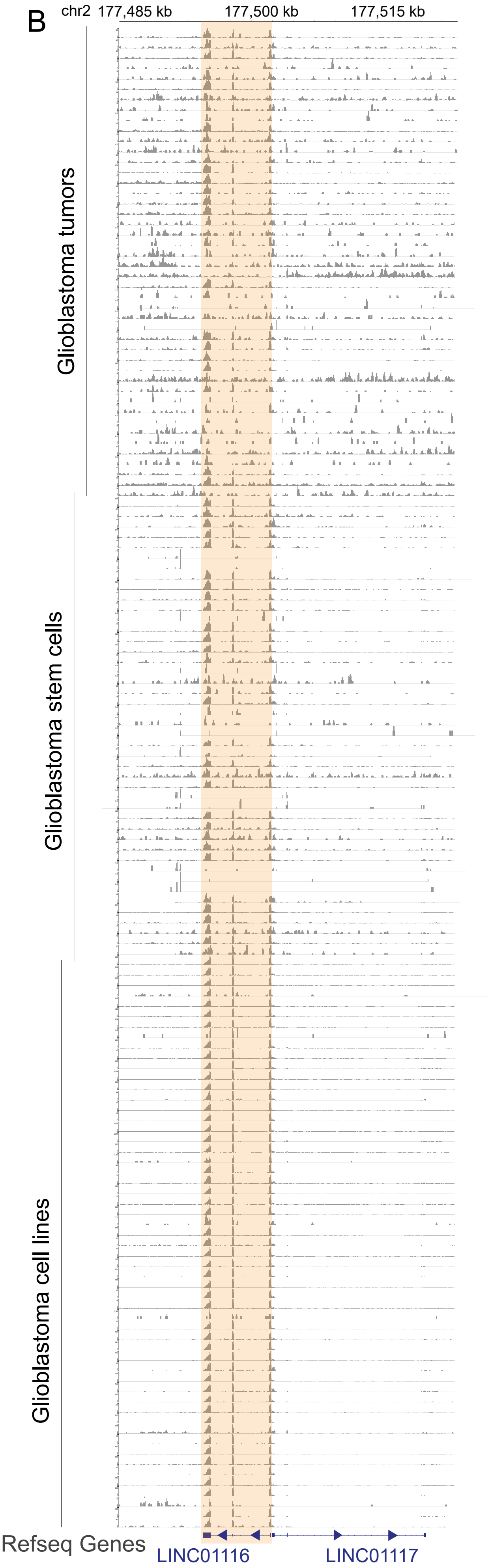


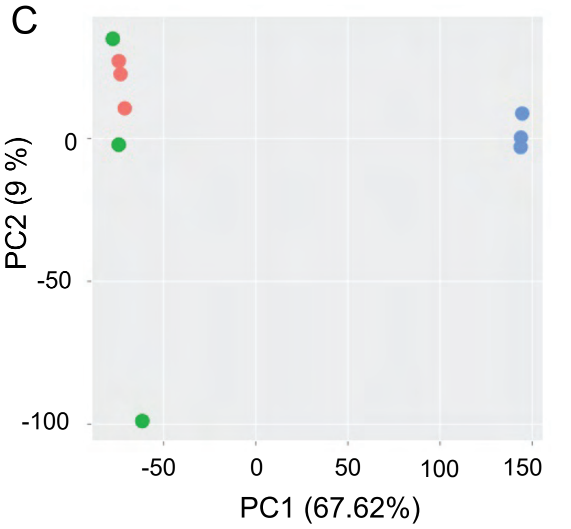


- Primary astrocytes
- Immortalized astrocytes - naive
- Immortalized astrocytes - transformed by HOXDeRNA

**Supplemental Figure 1. HOXDeRNA is not expressed in normal neuroglial cells, in contrast to glioma cells, and its activation globally alters transcriptional programs. Related to Figure 1.**
A-B. HOXDeRNA expression monitored by RNA-Seq in normal brain cells (GSE122701, GSE157461, GSE166847, GSE73721 and GSE119834 datasets) (A) and GBM tissues, GSCs (GSE119834), and glioma Cancer Cell Line Encyclopedia (Barretina et al., 2012) (B). HOXDeRNA transcript, composed of 3 exons, is expressed in all glioma samples but not in the normal cells of the brain.
C. Transcriptomic differences among primary astrocytes (n=3) and immortalized astrocytes naive or transformed by HOXDeRNA (n=3), visualized as PCA plot.

### Supplemental Figure 2

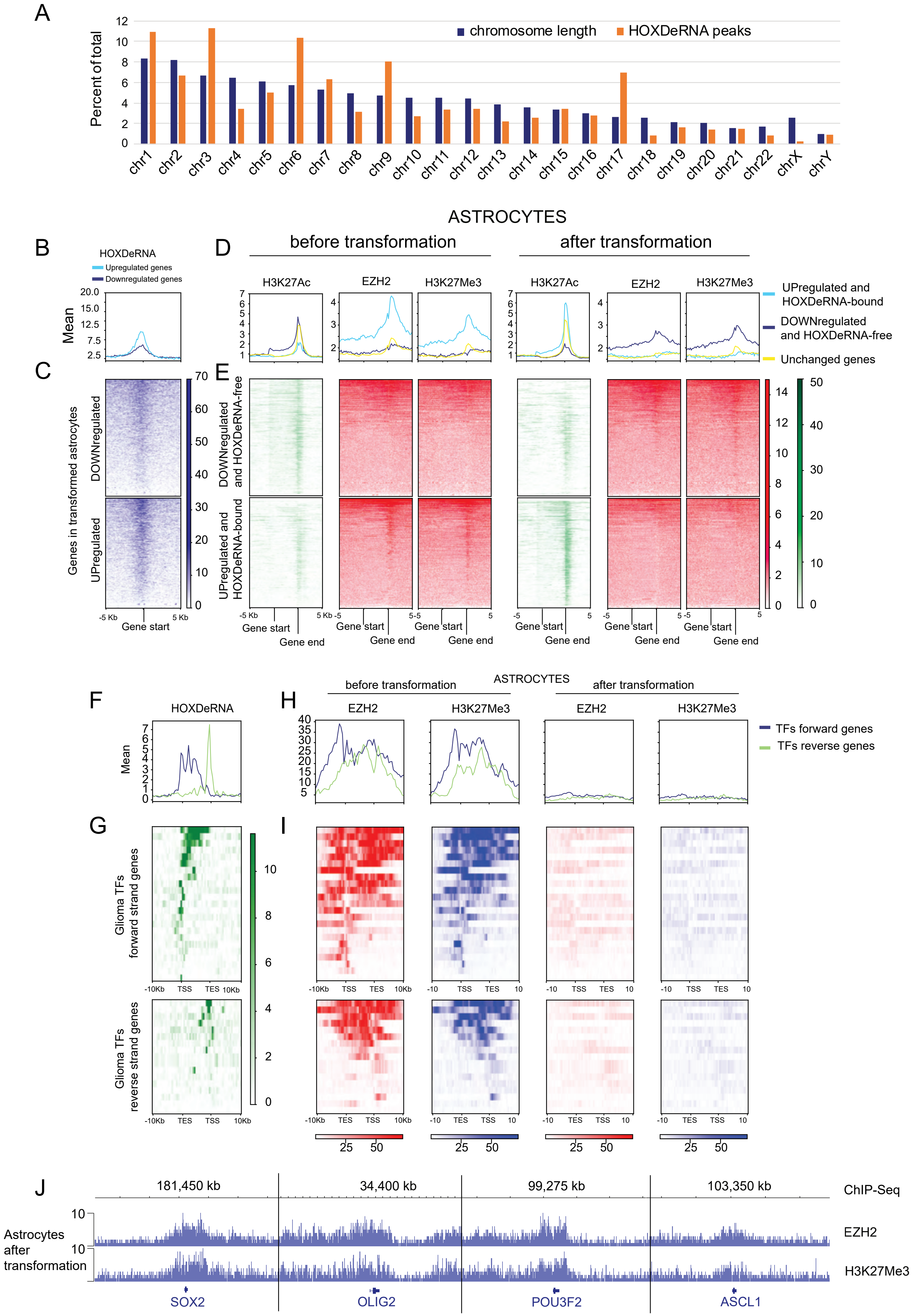

**Supplemental Figure 2. Genome-wide binding of HOXDeRNA is associated with exclusive PRC2 removal from the transformation-induced genes. Related to Figure 2.**

A. HOXDeRNA binds to all chromosomes. Relative chromosome length and number of HOXDeRNA peaks were calculated for each chromosome as percent of total.

B, C. ChIRP-seq raw read coverage signal, representing the HOXDeRNA binding at the TSS (+/- 5Kb) of the reverse strand of genes downregulated or upregulated after astrocyte transformation, is visualized as an average value for each group (B) or for each individual gene (C). The heatmap is accompanied by a colour scheme indicating the value of the raw read counts.

D, E. Epigenetic status of genes upregulated or downregulated in transformed astrocytes. H3K27Ac, H3K27Me3, and EZH2 ChIP-seq raw signals covering gene bodies of the negative strand (+/- 5Kb) were normalized to gene length and counted, followed by visualization of the average signal for 3 groups: genes upregulated after astrocytes transformation and bound by HOXDeRNA, downregulated after astrocyte transformation and HOXDeRNA-free, unchanged genes (D). Read coverage values for every individual gene body were visualized as heatmaps for negative strand genes, which are enriched in control or transformed astrocytes (E).

F, G. Genes encoding glioma master TFs are bound by HOXDeRNA after astrocyte transformation. HOXDeRNA ChIRP-Seq raw read signal was normalized to gene length and visualized as mean value (F) or individual values (G) for each TF gene body (+/- 10Kb) on forward and reverse strands. The colour scheme represents the values of the raw read counts.

H, I. PRC2 is removed from genes encoding glioma master TFs after astrocyte transformation. The EZH2 and H3K27Me3 raw read coverage signal was normalized to gene length and visualized as mean value (H) or individual values (I) for each gene body (+/- 10Kb) on both forward and reverse strands. The colour scheme represents the values of the raw read counts.

J. High-sensitivity H3K27Me3 and EZH2 ChIP-seq tracks in transformed astrocytes visualized for selected glioma master TF genes indicate incomplete removal of PRC2 activity in bulk chromatin after transformation. A different scale is presented in Fig. 2G bottom tracks to visualize relative levels in both non-transformed and transformed astrocytes.

### Supplemental Figure 3

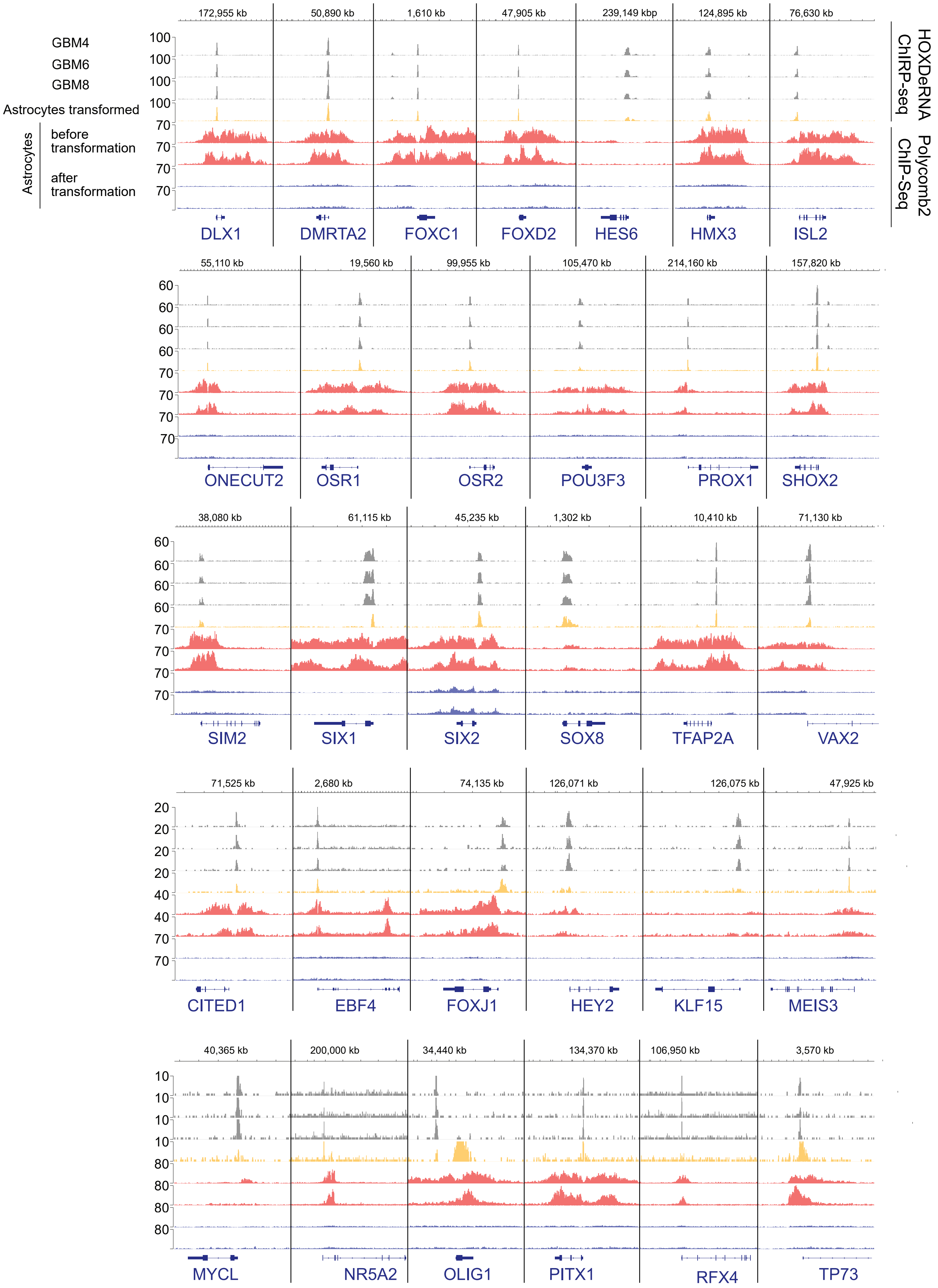

**Supplemental Figure 3. HOXDeRNA binds to and removes PRC2 repression from multiple glioma master TFs. Related to Figure 2.**

HOXDeRNA ChIRP-seq tracks in GSCs and transformed astrocytes, aligned with PRC2 (H3K27Me3, EZH2) ChIP-seq coverage, before and after astrocyte transformation, and visualized for glioma master TF genes.

### Supplemental Figure 4

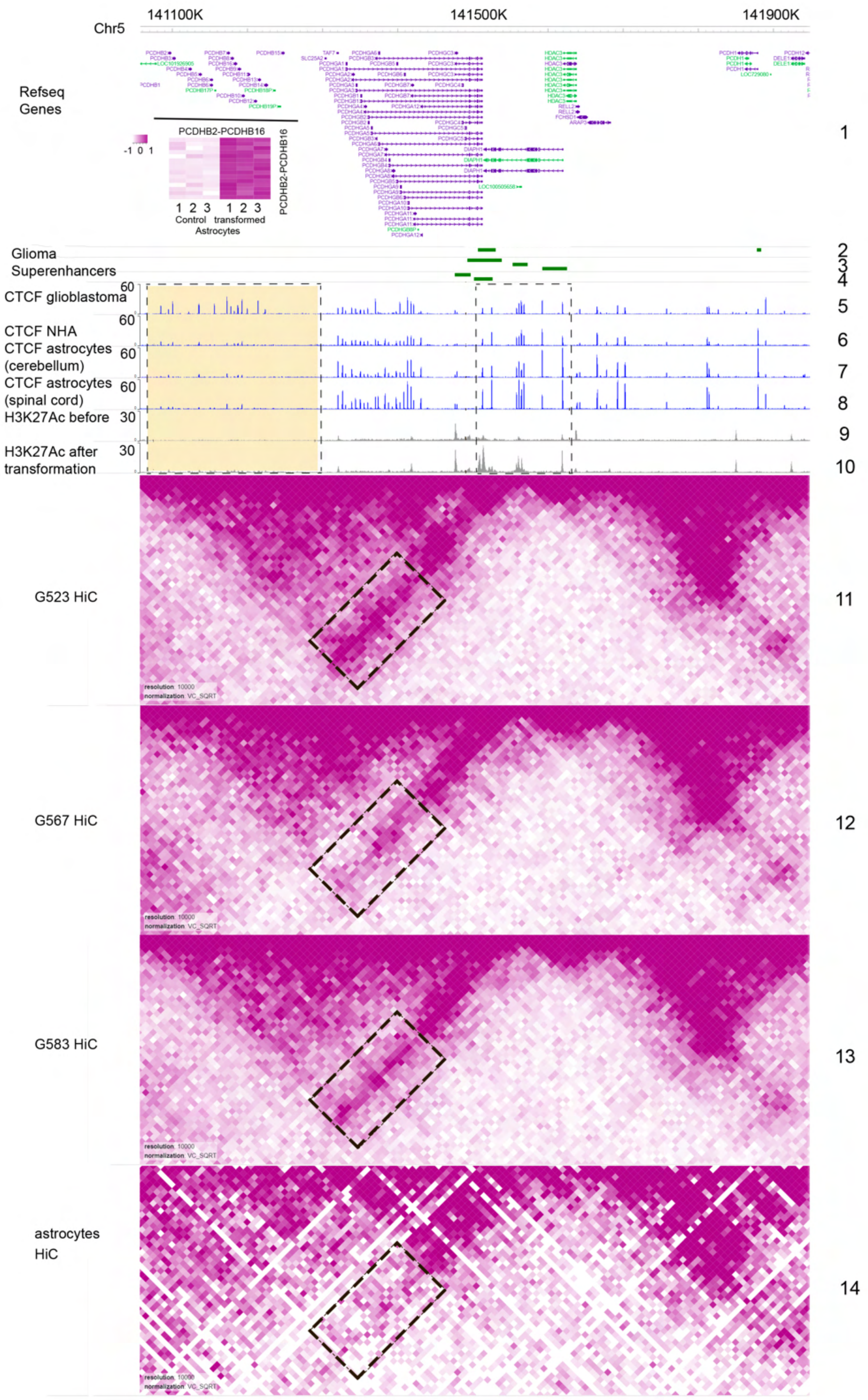

**Supplemental Figure 4. Integration of astrocytes RNAseq and ChIPseq data with ENCODE and GEO datasets suggests a mechanism of Protocadherin (PCDHB) family upregulation in GBM. Related to Figure 3.**

List of tracks from top to bottom: 1) Protocadherin genes annotated with Refseq. A heatmap visualizing PCDHB2-16 RNAseq-based expression in control and transformed astrocytes, transformed to z-scores; 2-4) SE tracks in three GCSs (GSE121601); 5-8) CTCF ChIP-seq tracks from ENCODE database for GBM (GSM822303) and three types of astrocytes (GSM733765, GSM1022662, GSM749696). Genomic region with differential CTCF binding is highlighted in yellow; 9-10) H3K27Ac coverage before and after astrocyte transformation shows activation of GBM specific SEs (dotted lines); 11-14) 4 HiC maps for GSCs G583, G567, G523 from (Johnston et al., 2019), available for download at https://wangftp.wustl.edu/hubs/johnston_gallo/), and astrocytes (GSE105194). A subset of HiC contacts representing SEs-protocadherins interactions are highlighted with dotted rectangles.

Supplemental Figure 5

A

| Position | Putative G4 forming sequences | G-Score | Name |
|---|---|---|---|
| 62 | GAGGGGAATGGGGAATCGGCAGGA | 33 | rG4-1 |
| 7241 | CAGGCTGGAGTGTGGTGGCA | 32 | rG4-2 |
| 3937 | CGGGCCGGGCGCGGTGGCT | 34 | rG4-3 |
| 3977 | TGGGAGGCCGAGGCGGGCGGAT | 34 | rG4-4 |
| 4111 | GGGAGGCTGAGGCAGGAGAATGGCGTGAACCCGG | 34 | rG4-5 |
| N/A | GAGAUGAATGAUGAATCGACAAGA | N/A | Control |

B

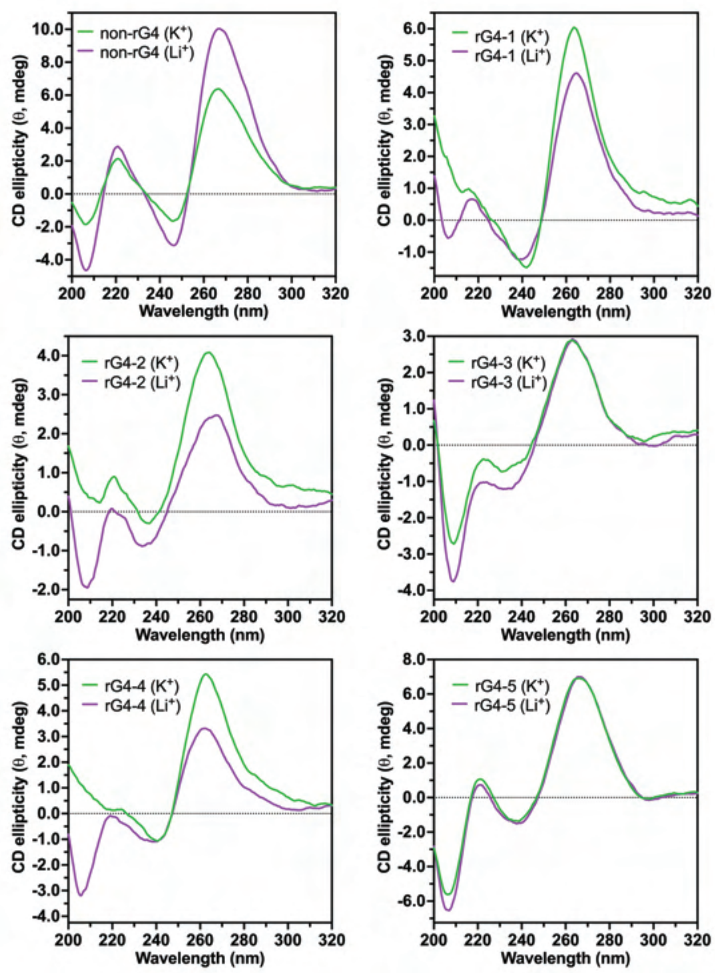


C

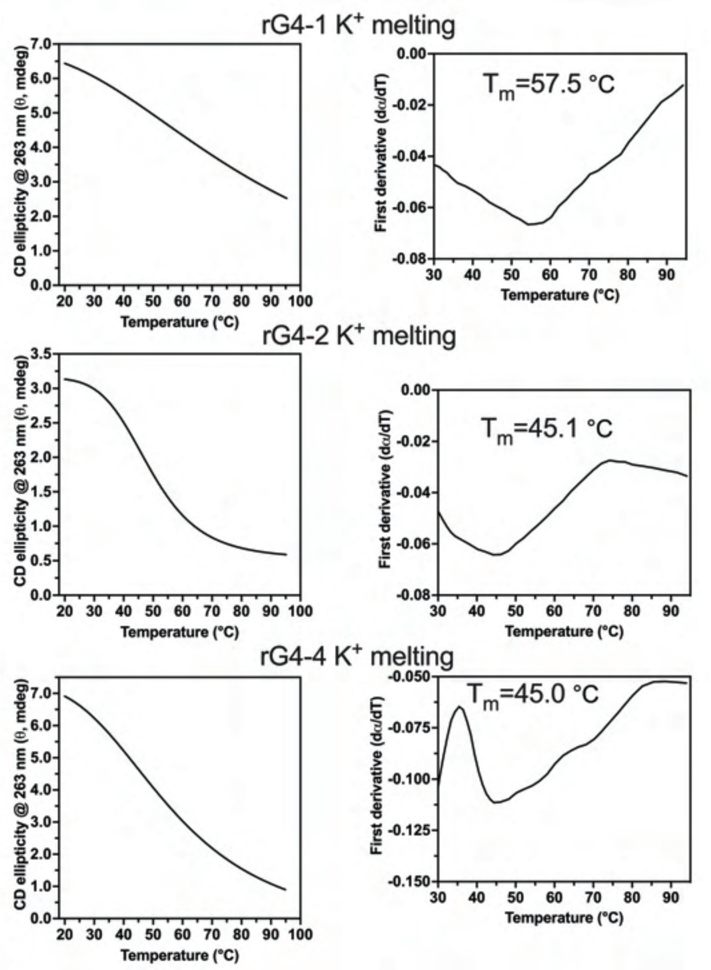


D

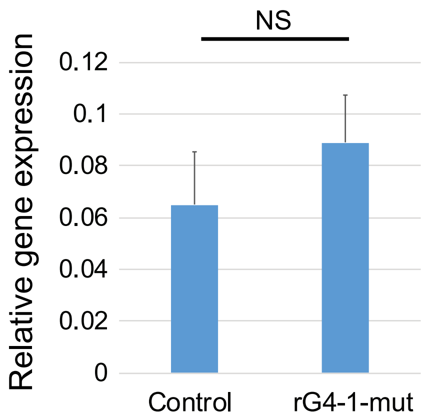

**Supplemental Figure 5. Characterization of rG4 candidates. Related to Figure 5.**
A. Five putative rG4 forming sequences detected with QGRS mapper tool. Search parameters: QGRS max length: 45, min G-Group size: 2, loop size: from 1 to 36.
B. CD spectra of five putative rG4 forming sequences (rG4-1, rG4-2, rG4-3, rG4-4, and rG4-5), along with one non-rG4 sequence. RG4-1, rG4-2, and rG4-4 exhibit common rG4 CD features (a higher intensity peak at 263 nm under K+ environment).
C. CD-melting curves indicating the stability of predicted rG4 structures. rG4-1 showed the highest melting temperature of 57.5 °C.
D. Base editing of rG4-1 does not alter HOXDeRNA expression levels in transformed astrocytes. qRT-PCR analysis, normalized to GAPDH mRNA levels (mean + SD, n=3).
